# Methods for addressing the protein-protein interaction between histone deacetylase 6 and ubiquitin

**DOI:** 10.1101/203372

**Authors:** Carolina dos S. Passos, Nathalie Deschamps, Yun Choi, Robert E. Cohen, Remo Perozzo, Alessandra Nurisso, Claudia A. Simões-Pires

## Abstract

Histone deacetylase 6 (HDAC6) is a cytoplasmic HDAC isoform able to remove acetyl groups from cellular substrates such as α-tubulin. In addition to the two deacetylase domains, HDAC6 has a C-terminal zinc-finger ubiquitin (Ub)-binding domain (ZnF-UBP) able to recognize free Ub. HDAC6-Ub interaction is thought to function in regulating the elimination of misfolded proteins during stress response through the aggresome pathway. Small molecules inhibiting deacetylation by HDAC6 were shown to reduce aggresomes, but the interplay between HDAC6 catalytic activity and Ub-binding function is not fully understood. Here we describe two methods to measure the HDAC6-Ub interaction *in vitro* using full-length HDAC6. Both methods were effective for screening inhibitors of the HDAC6-Ub protein-protein interaction independently of the catalytic activity. Our results suggest a potential role for the HDAC6 deacetylase domains in modulating HDAC6-Ub interaction. This new aspect of HDAC6 regulation can be targeted to address the roles of HDAC6-Ub interaction in normal and disease conditions.

---

Histone deacetylases (HDACs) are histone-modifying enzymes associated with several cell pathways including, of particular note, the transcriptional regulation of tumor repressor genes. Their extensive investigation as targets in drug discovery for cancer therapy has led to the development of small molecules that inhibit catalysis by different HDACs.^1^ Although transcriptional regulation is one of the mechanisms underlying the efficacy of HDAC inhibitors, part of the mode of action of pan-HDAC inhibitors has been associated with the catalytic inhibition of HDAC6, a cytosolic class IIb-type HDAC known to act on non-histone substrates.^2^ The role of HDAC6 in the clearance of misfolded proteins is of particular interest. Upon proteasome impairment, ubiquitin (Ub)-conjugated protein aggregates are recruited by HDAC6 as a compensatory pathway of elimination.^3, 4^ The combination of proteasome inhibitors with HDAC6 catalytic inhibitors has been shown to be effective in multiple myeloma; these results have encouraged the development of deacetylase inhibitors selective to HDAC6 for clinical applications.^3–5^

HDAC6 is unique among HDAC enzymes in that it possesses two catalytic deacetylase domains (CD1 and CD2) and a C-terminal zinc-finger Ub-binding domain (ZnF-UBP)^6^ (Fig. 1a). Strong evidence suggests that CD1, CD2, and ZnF-UBP domains all play a role in regulating aggresome formation and clearance by HDAC6.^7–9^ Nevertheless, the mechanisms on how catalytic inhibitors of HDAC6 might modulate the HDAC6-Ub interaction remained unclear until now. One hypothesis is that the catalytic inhibition would ultimately promote HDAC6 mislocalization and change the profile of HDAC6 protein-protein interactions (PPIs). Another hypothesis is that the binding of small molecules to the catalytic domains of HDAC6 would promote conformational changes that disrupt the interaction between the HDAC6 ZnF-UBP domain and Ub. The latter scenario can be assessed most directly by use of full-length HDAC6 in cell-free protein interaction assays.

**Fig. 1.**
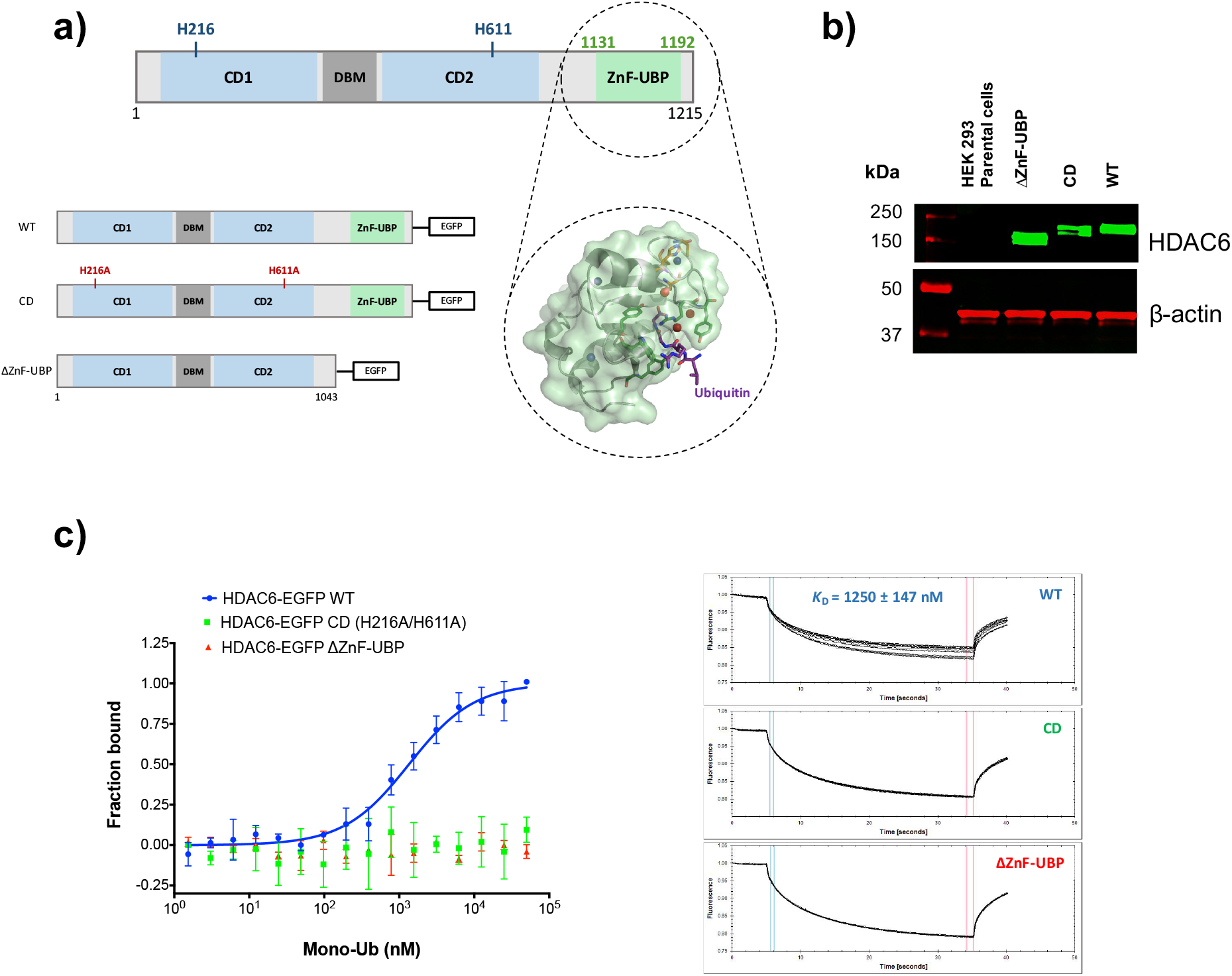
Overexpression of HDAC6-EGFP fusion proteins in HEK 293 cells and their interaction with recombinant mono-Ub measured by MST. **a)** Scheme of the full-length HDAC6 depicting its catalytic domains (CD1, CD2) and the zinc-finger ubiquitin binding domain (ZnF-UBP) able to recognize the C-terminal GG motif of Ubiquitin (Ub). Detail: molecular surface of HDAC6 ZnF-UBP in complex with Ub C-terminus RLRGG (black ball & sticks). Image generated with MOE 2015.10 from PDB: 3GV4. **b)** Western blot analysis of cell lysates from HEK 293 cells overexpressing HDAC6-EGFP fusion proteins: wild-type (HDAC6-EGFP WT), catalytic deficient (HDAC6-EGFP CD), and ZnF-UBP deleted (HDAC6-EGFP ΔZnF-UBP). **c)** MST titration binding experiments showing the interaction between HDAC6-EGFP WT fusion protein and mono-Ub, and the absence of interaction between HDAC6-EGFP CD / ΔZnF-UBP and mono-Ub.

Up to now, a variety of binding assays using the isolated HDAC6 ZnF-UBP domain have been employed to evaluate the HDAC6-Ub interaction;^10, 11^ however, these are not suitable for investigating potential effects of the catalytic deacetylase domains on the HDAC6-Ub interaction. Here we describe *in vitro* methods that we have developed to (1) evaluate the interplay between catalytic deacetylase domains and the ZnF-UBP Ub-binding domain in HDAC6, and to (2) identify small molecules that interact with the ZnF-UBP and inhibit the HDAC6-Ub interaction.

## Results and discussion

Using microscale thermophoresis (MST) and human full-length HDAC6, we developed a binding assay based to detect the specific binding of proteins and small drug-like molecules to the ZnF-UBP domain of HDAC6. MST is a fluorescence-based technique with a wide range of applications in the study of biomolecular interactions.^12^ Binding of molecules to a fluorescent protein will impact its thermophoretic behavior due to changes in the protein’s hydration shell, size, or charge upon binding. In addition, when ligand binding occurs close to the fluorophore environment, MST can be used in the temperature jump (T-Jump) mode to evaluate changes triggered by binding. A particular advantage of the MST approach is that the assays can use cell lysates directly with proteins expressed in cells as fusions to fluorescent proteins, thereby avoiding subsequent purification, immobilization, or fluorescent-labeling steps that could alter protein folding and activity.^12^ Indeed, we have previously observed loss of binding activity with the full-length HDAC6 while attempting amine-coupling reactions for fluorescent-labeling (in MST approach) or surface-immobilization on surface plasmon resonance (SPR) (data not shown).

To provide suitable fluorescent interaction partners, HEK 293 cells were transiently transfected with full-length HDAC6-EGFP constructs to obtain either the wild type (HDAC6-EGFP WT), the catalytic deficient (HDAC6-EGFP CD), or the ZnF-UBP deleted (HDAC6-EGFP ΔZnF-UBP) proteins (Fig. 1a-b). In each of these constructs, EGFP was fused to the HDAC6 C-terminus, which is located near ZnF-UBP. MST experiments then were conducted with the whole-cell lysates prepared as described in the Methods to increase HDAC6 folding stability and maintain its catalytic activity at the same time (Fig. S1). HDAC6-Ub binding curves using lysates containing HDAC6-EGFP WT, CD, and ΔZnF-UBP proteins were obtained by evaluating thermophoresis + T-Jump (Fig. 1c). The dissociation constant (*K*_D_) for the interaction between mono-Ub with the full-length HDAC6-EGFP WT was determined (*K*_D_ = 1250 ± 147 nM). Experiments with different dilutions of lysates from cells transfected with HDAC6-EGFP WT showed similar *K*_D_ values for the HDAC-Ub interaction suggesting no significant interference by endogenous free Ub, Ub-chains, or Ub-like proteins (Fig. S2). The *K*_D_ for His_6_-tagged Ub binding to HDAC6, which shows ~15-fold higher affinity than untagged monoUb, was in accordance with previously published ITC data using the ZnF-UBP domain alone^13^ (Fig. S3). HDAC6 mutations H216A/H611A (HDAC6-EGFP CD), which lead to complete catalytic impairment, resulted in the loss of the HDAC6-Ub interaction (Fig. 1c). This suggested that structural changes in the catalytic domains may affect the Ub-binding activity of HDAC6. Binding studies with Ub-chains and Ub-like proteins (**Fig. S4-5**) also were done to confirm the versatility of the MST approach for determining the affinity constants of HDAC6 PPIs *in vitro*.

An important application of the method is the screening of small molecules able to interact with HDAC6 residues involved in Ub recognition and that potentially can act as direct PPI inhibitors. To explore this application, a structure based virtual screen (SBVS) was first conducted to identify compounds potentially targeting the HDAC6 ZnF-UBP.^10, 14^ From a chemical library of 197,477 compounds (www.specs.net, SC_specs_10 mg_May2013), 40 potential hits (Table S1) were selected and tested by MST for their binding to HDAC6-EGFP WT using thermophoresis + T-Jump (Fig. 2a). This measurement provided a suitable Z-factor (0.6) for medium-throughput screening and included the contribution of the ZnF-UBP binding via T-Jump. To rank the screened molecules, we used 50 µM mono-Ub as a fraction bound control in the screening experiment. Normalized fraction-bound values were determined by comparing the MST curves of the tested compounds to those obtained for the unbound (lysate alone) and bound (lysates pre-incubated with 50 µM mono-Ub) controls. A normalized fraction bound of 0.5 was considered as the threshold resulting in the selection of one hit, compound **16**, which had its potential binding mode predicted by molecular docking (Fig. 2b). Unfortunately, MST experiments to determine the *K*_D_ for the binding between compound **16** and HDAC6-EGFP WT could not be performed because of the strong intrinsic fluorescence displayed by the ligand at concentrations higher than 100 µM.

**Fig. 2.**
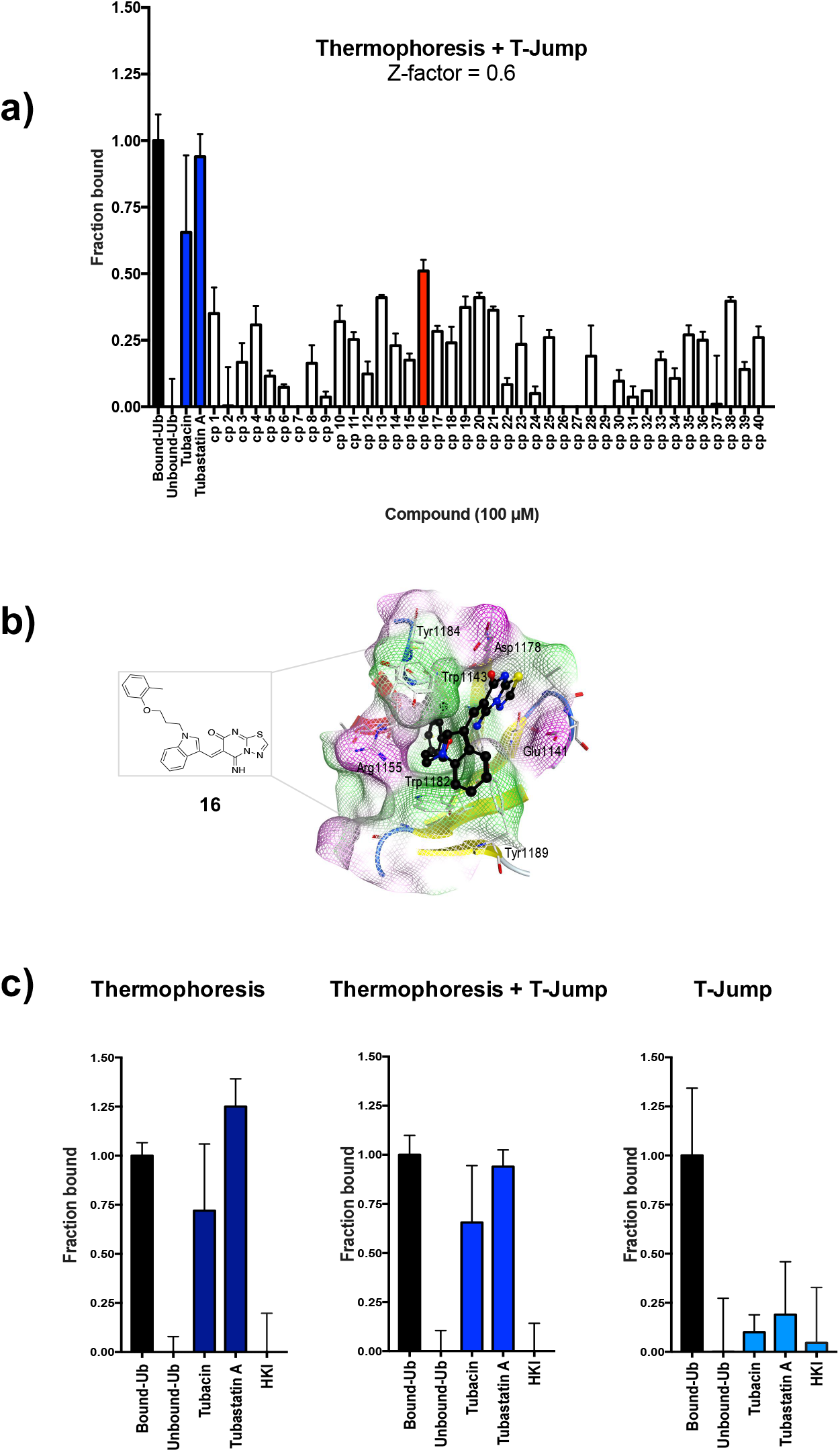
Microscale thermophoresis (MST) applied for the screening of HDAC6 ligands: detection of binding on both catalytic and ZnF-UBP domains. **a)** Selected compounds were screened at 100 μM. Fraction bound values were obtained by normalizing the amplitude of thermophoresis + T-Jump values (between 100 and 0 µM compound) over the amplitude values between bound (50 μM mono-Ub) and unbound (0 μM mono-Ub) controls. **b)** Best ranked docking pose for compound **16** in complex with HDAC6 ZnF-UBP. Compound **16** is represented as balls & sticks. 2D structure is also depicted. The molecular surface around the ligand is color-coded based on lipophilicity (green: hydrophobic; purple: hydrophilic). Key amino acids surrounding **16** are labelled in black. Image was generated with MOE 2015.10. **c)** Comparison of MST measurements with HDAC6-selective deacetylase inhibitors tested at 100 µM: T-Jump analysis detects preferentially binding close to the fluorophore (EGFP is fused to the C-terminus of HDAC6) and to a lesser extent the binding of catalytic inhibitors.

In addition to the positive binding control mono-Ub (ZnF-UBP-site positive, catalytic-site negative), the selective HDAC6 inhibitor tubastatin A (ZnF-UBP-site negative, catalytic-site positive binding control) and a HDAC class I-selective hydroxyl ketone inhibitor (HKI; ZnF-UBP-site negative, catalytic-site negative binding control, Fig. S6)^15^ were probed. Binding of HDAC6 catalytic inhibitors to the full-length HDAC6 could be detected in both thermophoresis alone and thermophoresis + T-Jump experiments (Fig. 2c), but only at a concentration (100 µM) well above their nanomolar catalytic IC_50_. Moreover, their T-Jump values alone (Fig. 2d) pointed to an absence of binding close to the fluorophore (i.e., near the ZnF-UBP), corroborating the use of MST thermophoresis + T-Jump for detecting binding to the HDAC6 ZnF-UBP domain.

Looking for small molecules able to bind the ZnF-UBP will not necessarily provide compounds able to disrupt the HDAC6-Ub PPI. In the interaction surface between two proteins, binding pockets able to accommodate small molecules (the potential ‘hotspots’) are much smaller than the protein-protein interface.^16, 17^ To check the ability of compound **16** to inhibit the HDAC6-Ub PPI specifically, and to confirm the screening results, an ELISA-based PPI competition assay was further developed and conducted with 40 compounds from the SBVS hits. In this assay, His_6_-tagged mono-Ub was immobilized on Nickel-coated 96-well plates. Tag-free mono-Ub (positive competition control) or tested compounds were incubated with recombinant HDAC6 WT and then added to the plates. The competition was evaluated by calculating the fraction of HDAC6 bound to immobilized His-tag mono-Ub, following recognition of the His_6_-Ub-HDAC6 complex by an anti-HDAC6 primary antibody (Fig. 3a). Concentration-response competition curves were first determined for controls titrated with tag-free mono-Ub (Fig. 3b). Then, the screening was conducted with the SBVS hits (Fig. 3c). Compound **16** was confirmed to disrupt the HDAC6-Ub PPI. Interestingly, tubacin and tubastatin A also inhibited the PPI (Fig. 3c). Such inhibition corroborates the MST results observed with the catalytic deficient HDAC6. Taken together, these results confirm for the first time that changes in the HDAC6 catalytic domains directly inhibit the HDAC6-Ub PPI, most likely via an allosteric-like mechanism. This was also observed, albeit to a lesser extent, with the pan-HDAC inhibitor SAHA. In contrast, class I selective inhibitors such as CI-994 and HKI had no or little effect on the Ub interaction with HDAC6 (Fig. 3c).

**Fig. 3.**
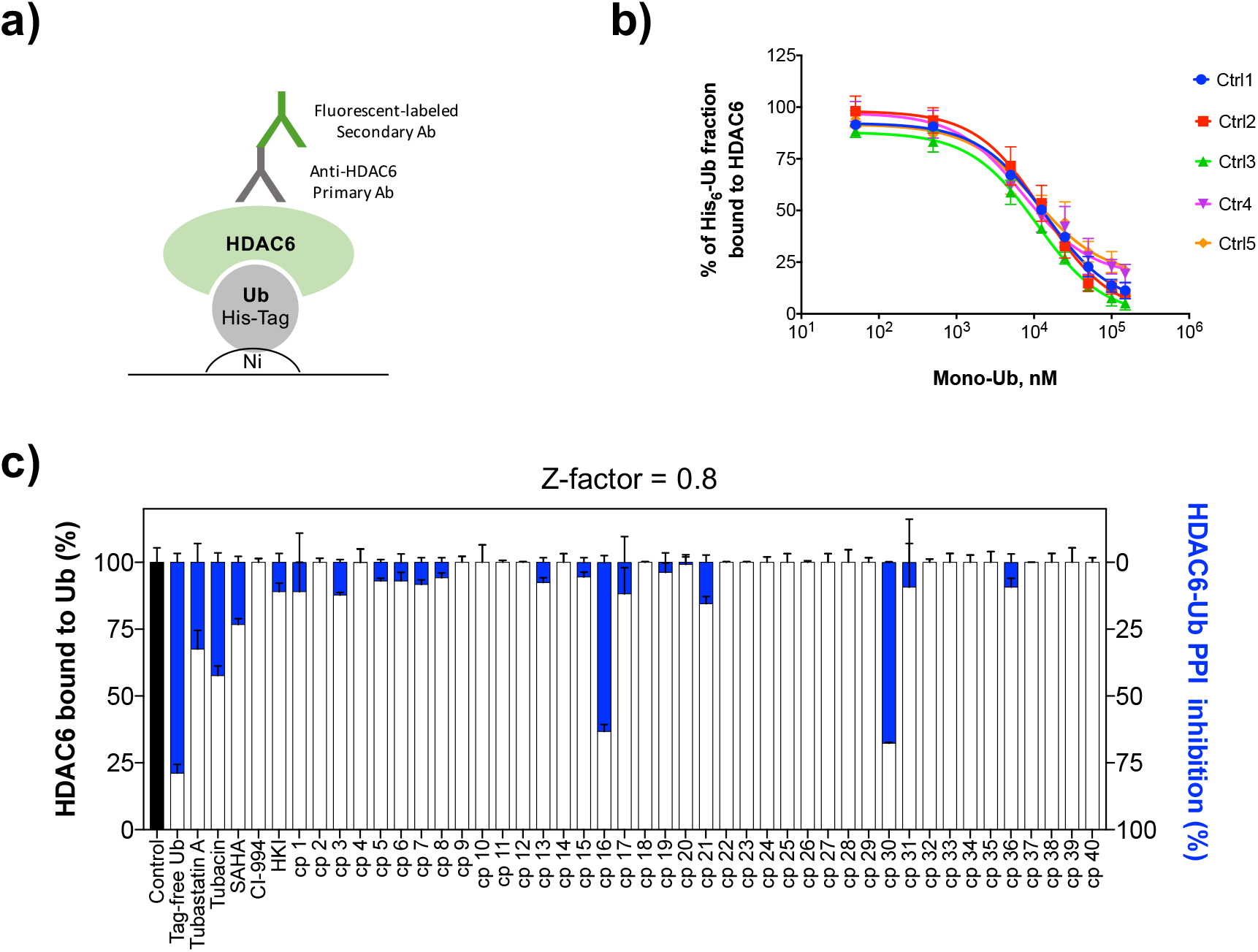
ELISA-based method for measuring the inhibition of the protein-protein interaction (PPI) between HDAC6 and Ub. **a)** His_6_-Ub was immobilized on Nickel (Ni) coated 96-wells plates and let to interact with recombinant HDAC6 WT. After washing steps to remove unbound proteins and ligands, plates were subjected to sequential incubations and washing with an anti-HDAC6 antibody and a fluorescent labeled secondary antibody. HDAC6 interaction with immobilized His_6_-Ub was detected by fluorescence measurements. **b)** Control experiments were performed to evaluated whether tag free mono-Ub could compete with immobilized His_6_-Ub for its binding to HDAC6. Competition curves from five independent experiments showed that pre-incubation of HDAC6 with increasing concentrations of tag free mono-Ub reduced the His_6_-Ub-HDAC6 complex formation in a concentration-dependent fashion (IC_50_ = 11.2 ± 2.7 μM). **c)** Screening of 40 selected SPECS compounds for HDAC6-Ub PPI inhibition by the ELISA-based method.

Compound **30** appeared as a second hit in the ELISA-based PPI competition assay, despite the absence interaction observable via the MST screening. Given that compounds binding at the catalytic domains, such as tubastatin A and tubacin, were shown to affect HDAC6-Ub interaction, hits **16** and **30** (Fig. S6) were tested for effects on HDAC6 deacetylase activity. No catalytic inhibition was observed in those assays, but, a 2-fold *increase* in substrate deacetylation was seen for compound **30** (Table S2). While the effect of compound **30** is unclear, it does not exclude its interaction with potential allosteric binding sites of HDAC6, able to modulate both its catalytic and its Ub-binding functions.

The MST assay using lysates of cells expressing full-length HDAC6-EGFP fusion proteins allowed investigation of the HDAC6-Ub interactions showing, for the first time, the role of HDAC6 catalytic inhibition on HDAC6-Ub PPI. Our results show that, in combination with an ELISA-based PPI assay, the MST method has the potential to identify HDAC6-Ub PPI inhibitors that can be uncoupled from effects on HDAC6 catalysis. Such compounds could be used as probes to explore mechanisms involved in aggresome regulation by HDAC6, which are of interest in cancer, viral infection, neurological diseases, and immune disorders.^1, 18^ By combining the MST assay with the ELISA-based PPI competition assay, we identified the first HDAC6-Ub PPI inhibitor (**16**). Overall, the approach developed here can be used to measure the interaction of the ZnF-UBP with its molecular partners, from small molecules to proteins, in the context of the full-length and physiologically relevant form of HDAC6.

## Methods

### 1. HEK 293 cell cultures, transfection, and cell lysates

HEK 293 cells (ATCC^®^ CRL-1573™) were cultured in Dulbecco’s Modified Eagle Medium (DMEM) with high glucose (Gibco™, ThermoScientific # 41965120), supplemented with 10% fetal bovine serum, 100 units/mL penicillin, 100 µg/mL streptomycin, and 1% non-essential amino acids. Cultures were kept in 75 cm^2^ culture flasks (NUNC, Thermo Scientific) at 37 ºC in a 5% CO2 incubator (HERAcell, Heraeus). Cells were transfected with pEGFP.N1-HDAC6, pEGFP.N1-HDAC6 DC, or pEGFP.N1-HDAC6.delta ZnF-UBP for 48 h. To prepare lysates, medium was removed from culture flasks and cells were washed twice with 10 mL ice cold Dubelcco’s Phosphate Bufered Saline (DPBS, Gibco™, ThermoScientific # 14190144). A total of 500 μL of ice cold radioimmunoprecipitation assay buffer (RIPA buffer, Sigma-Aldrich # R0278) containing protease inhibitor cocktail (SIGMA*FAST*™, Sigma-Aldrich # S8820) was added to each flask, and cells were scraped to provide cell suspensions in lysis buffer. The cell suspensions were transferred into conical tubes, vortexed for 2 s and kept on ice for 20 min. Tubes were then vortexed again (2 s) and centrifuged at 13000 x g for 10 min at 4 °C. The supernatants were transferred to new tubes as whole-cell lysates. For each lysate, an aliquot was immediately used for EGFP determination. EGFP concentrations in the lysates were measured by fluorescence in the Monolith NT.115 Instrument (MO-G008, LED power 60%, excitation 460-480 nm, and emission 515-530 nm) using recombinant GFP (rGFP) as a standard (Roche Diagnostic AG # 11814524001). Lysates were stored at −80 °C, divided into aliquots (to avoid freeze-thaw cycles), and thawed on ice prior to MST experiments. EGFP concentration and binding affinities were shown to be stable across aliquots.

### 2. Antibodies, proteins, plasmids, and reagents

Antibodies: HDAC6 (D21B10) Rabbit mAb (# 7612), β-actin (8H10D10) Mouse mAb (# 3700), Anti-rabbit IgG (H+L) (DyLight™ 800 4X PEG Conjugate) (# 5151), Anti-mouse IgG (H+L) (DyLight™ 680 Conjugate) (#5470) (all from Cell Signaling Technology, Danvers, MA, USA), Anti-Ubiquitin antibody [EPR8830] Rabbit mAb (# 134953, Abcam, Cambridge, UK). Goat monoclonal anti-rabbit IgG-FITC (#SC-2012) was from Santa Cruz Biotechnology. Proteins: Ubiquitin, human, recombinant (# BML-UW-0280), Ubiquitin, human, recombinant, His-tag (BML-UW8610), NEDD8, human, recombinant, His-tag (BML-UW9225), ISG15, human, recombinant, His-tag (BML-UW9335) (all from Enzo Life Sciences, Inc.), GST-HDAC6, human, recombinant (Sigma-Aldrich, # SRP0108).

Plasmids: pEGFP.N1-HDAC6, pEGFP.N1-HDAC6.DC, and pEGFP.N1-HDAC6.delta ZnF-UBP were a gift from Tso-Pang Yao (Addgene plasmids # 36188, 36189, and 36190, respectively).^19^

Reagents: Fluor de Lys^®^ SIRT1 substrate (BML-KI177), Fluor de Lys^®^ developer II (BML-KI176) (all from Enzo Life Sciences, Inc.), trichostatin A (Sigma-Aldrich, #T8552), tubastatin A (Apex Bio, #A410), tubacin (Sigma-Aldrich, # SML0065), SAHA (Sigma-Aldrich, # SML0061), CI-994 (Sigma-Aldrich, # C0621). The class I-selective hydroxyl ketone compound was a gift from Dr. Yung-Sing Wong.^15^ All compounds screened for ZnF-UBP-Ub modulation were purchased from the Specs company (http://www.specs.net).

### 3. Ubiquitin chains

K48- and K63-linked ubiquitin chains were synthesized as previously described^20^, using linkage-specific enzymes to catalyze the reactions. The mammalian enzyme E2-25K was used in the synthesis of K48-linked chains while the yeast complex Mms2/Ubc13 was used to synthesize K63-linked chains. To generate di-Ub chains, the Ub moiety forming the proximal end carried unmodified K48 or K63 and an extra C-terminal residue (D77). The Ub moiety forming the distal end carried unmodified C-terminal G76 and K48C or K63C mutants. After synthesis of Ub-chains, the C-terminal D77 residue in the proximal Ub was removed through reaction with a recombinant yeast ubiquitin C-terminal hydrolase (Yuh1). Polyubiquitin chains were extended to form K48- or K63-linked tetra-Ub by successive rounds of deblocking and conjugation.

### 4. SDS-page and immunoblotting

Lysates of HEK 293 cells overexpressing the EGFP fusion proteins were diluted 1:1 with sample buffer (240 mM Tris/HCl pH 6.8 containing 0.04% bromophenol blue, 40% glycerol with or without 200 mM DTT), heated at 70 ºC for 10 min, and separated on 8% polyacrylamide gel. Before binding experiments, lysates of HEK 293 cells were incubated with mono-Ub or Ub-chains (K48-linked di- and tetra-Ub, or K63-linked di- and tetra-Ub) and monitored by SDS-page and immunoblotting to test for chain degradation by deubiquitinating enzymes (DUBs) possibly present in the cell lysates. Lysates (2- to 4-fold dilution) were pre-incubated with 4 µM ubiquitin aldehyde (Ubal; K48- and K63-linked Ub chains) and 4 mM 1,10-phenanthroline (1,10-phen; K63-linked Ub chains only) for 15 min at 37 ºC, followed by incubation with K63- or K48-linked Ub chains, for 10 min at 37 ºC. Samples were then diluted 1:1 with sample buffer, incubated for 20 min at room temperature, and separated on 15% polyacrylamide gel.

For immunoblotting, proteins were transferred to PVDF membranes, and the membranes were subsequently blocked with TBS containing dried-milk powder (5%) and Tween^®^-20 (0.05%) for 1 h at 4 ºC. Membranes were then incubated with primary antibodies diluted 1:1000 (anti-HDAC6 and anti-Ub rabbit monoclonal) or 1:5000 (anti-β-actin mouse monoclonal) in the blocking buffer overnight at 4 ºC. Membranes were washed three times with TBS containing Tween^®^-20 (0.05%) and incubated with fluorescent secondary antibodies diluted 1:10000 in blocking buffer for 1 h at room temperature. Membranes were washed as previously described, and the results were visualized with an Odyssey infrared imaging system (LI-COR Biosciences).

### 5. Structure-based virtual screening (SBVS)

Specs compounds were prepared for SBVS as previously described.^21^ Briefly, the compounds were downloaded from the Specs website (www.specs.net, SC_specs_10 mg_May2013) and protonated at pH 7.4 using the Protonate 3D tool of MOE 2013.08 (Chemical Computing Group CCG, Montreal, Canada), followed by energy minimization (MOE, MMFF94x force field, R-field equation for solvation, and 0.1 kcal/mol/Å2 as RMS gradient cut-off). Raccoon v1.0^22^ was used to convert the ligand input files to the .pdbqt format. Atomic coordinates for HDAC6 ZnF-UBP (PDB 3GV4) were retrieved from the Protein Data Bank (http://www.wwpdb.org/). Protein structure was prepared using the AutoDock Tools (ADT) module of MGLTools v1.5.6.^23^ The co-crystallized ubiquitin C-terminal peptide RLRGG and water molecules were removed from the protein structure, followed by the addition of Gasteiger charges and non-polar hydrogens. The binding site was centered on the carboxyl group of the C-terminal glycine in the co-crystallized peptide RLRGG. Box dimensions were 20 × 18 = 20 Å. Docking calculations were performed with AutoDock Vina^24^ using default parameters. 20 binding poses were generated for each compound. The best-ranked compounds with scores < −8.5 kcal/mol were further filtered for solubility (> 100 µM) and logP (< 5.0), as predicted with VolSurf + version 1.0.7,^25^ resulting in 609 compounds that were visually inspected to ensure proper fit to the Ub binding pocket. 80 ligands were selected for re-docking in HDAC6 ZnF-UBP using GOLD version 5.2 (CCDC, Cambridge, UK). The binding site was defined to be within 6 Å of the co-crystallized peptide. 100 docking poses were generated for each Specs compound by using 100,000 GOLD Genetic Algorithm interactions (Preset option) and ranked according to the ChemPLP score. Finally, 40 compounds with a ChemPLP score higher than 70 were selected for *in vitro* binding experiments.

### 6. Microscale thermophoresis (MST)

Solutions and serial dilutions of non-tagged mono-Ub, K48-linked di-Ub, and K63-linked di- and tetra-Ub were prepared in 50 mM Tris/HCl buffer pH 8.0 containing 150 mM NaCl; solutions of K48-linked tetra-Ub (K48C mutation in the distal Ub moiety) were in this same buffer supplemented with 5 mM DTT. Solutions and serial dilutions of His_6_-tagged mono-Ub, NEDD8, and ISG15 were prepared in 20 mM Hepes pH 8.0 containing 50 mM NaCl and 1 mM DTT. Lysates of HEK 293 cells overexpressing wild type (WT), catalytic deficient (CD), or ∆ZnF-UBP forms of EGFP-HDAC6 were used as the source of labeled fluorescent proteins for MST experiments. In addition, control experiments were done with non-tagged mono-Ub labeled with the fluorescent dye NT-647 (Monolith NT™ Protein Labeling Kit RED-NHS, NanoTemper Technologies GmbH, München, Germany) and purified GST-HDAC6 WT (Fig. S7). Protein concentrations of unlabeled mono-Ub, Ub chains, Ub-like proteins, and GST-HDAC6 stock solutions were measured using the Qubit^®^ Protein Assay (ThermoFisher Scientific, # Q33211).

Protein-protein interactions between HDAC6 and mono-Ub were first evaluated by MST with the isolated protein partners: fluorescent-labeled mono-Ub and GST-HDAC6. To determine the dissociation constants, 10 µL of 132 nM labeled mono-Ub (50 mM Tris/HCl pH 8.0, 137 mM NaCl, 2.7 mM KCl, 1 mM MgCl2, 1 mg/mL BSA) were mixed with 10 µL of unlabeled GST-HDAC6 WT in concentrations ranging from 0.55 to 8931 nM (serial dilutions in the same buffer plus 10% glycerol). After 15 min at room temperature, samples were loaded into MST ‘premium’ coated capillaries (NanoTemper Technologies, # MO-K005), and the thermophoresis behavior of each was measured (excitation 605-645 nm; emission 680-685 nm) using the Monolith NT.115 instrument (NanoTemper Technologies). Measurements were performed at room temperature for 30 s at 60% LED power and 40% infrared laser power (Fig. S7).

For MST experiments with tag-free mono-Ub (Fig. 1) and Ub chains (Fig. S4), lysates of cells expressing EGFP fusion proteins were diluted 2 to 4-fold in lysis buffer and pre-incubated with deubiquitination enzyme inhibitors (i.e., 4 µM Ubal alone or plus 4 mM 1,10-phenanthroline) for 15 min at 37 ºC as described above. Then, 10 µL of lysate was mixed with 10 µL of unlabeled mono-Ub or polyUb chains, incubated for 10 min at 37 ºC, loaded into MST premium coated capillaries, and thermophoresis was measured (excitation 460-480 nm; emission 515-530 nm) using the Monolith NT.115. Lysates were tested in final dilutions between 4- and 8-fold (final EGFP concentrations of approximately 25 nM). Each measurement was performed at room temperature for 30 s at 95% LED power and 40% infrared laser power. Experiments with His_6_-tagged mono-Ub and Ub-like proteins were performed according to the same protocol without pre-incubation with Ubal and 1,10-phenanthroline; unlabeled proteins ranged from 0.70 to 60,000 nM. Dissociation constants were determined from fits to the MST data using the law of mass action Kd formula available as a Monolith NT.115 data analysis tool. Fraction-bound values were calculated for each concentration by subtracting the baseline in each experiment and dividing by the amplitude; fraction-bound values were then fitted in GraphPad^®^ Prism 7.0 nonlinear regression equation “log (agonist) vs. response (three parameters)” from the Prism equation library.

The optimized MST assay was used to test the 40 compounds selected by SBVS for their binding to HDAC6. Briefly, 100 µM of the tested compounds were pre-incubated (10 min, 37 °C) with lysates of HEK293 cells expressing HDAC6-EGFP WT. MST measurements were performed at room temperature for 30 s at 95% LED power and 40% infrared laser power. MST curves for the tested compounds were compared to those obtained for unbound (lysate alone) and bound (lysates pre-incubated with 50 µM ubiquitin) controls. Fraction-bound values were determined from triplicate measurements by subtracting the baseline of the unbound control and dividing by the amplitude of the bound control (50 µM ubiquitin).

### 7. HDAC6 catalytic assay

*In vitro* HDAC6 activity was detected by measuring the fluorophore released from the Fluor de Lys^®^ acetylated substrate by the deacetylase activity. Trichostatin A (a pan-HDAC inhibitor) was used as a control for HDAC6 inhibition. Stock solutions of trichostatin A and tested molecules in DMSO were diluted in HDAC6 assay buffer (50 mM Tris/HCl pH 8.0, 137 mM NaCl, 2.7 mM KCl, 1 mM MgCl2, 1 mg/mL BSA). The final DMSO concentration was kept at 2%. Assays were conducted in duplicate in 96-well plates. Compounds were added to GST-HDAC6 solution (50 U/reaction) and the reaction was initiated by adding 50 μM acetylated substrate, followed by incubation at 37 °C for 1 h. Enzymatic reactions were stopped by adding 1 μM trichostatin A. A peptidase reagent (Fluor de Lys^®^ developer II) was added) and incubated for 3 h at RT to release the fluorophore from the deacetylated product. The resulting fluorescence intensity was measured at 360 nm excitation and 460 nm emission, using a FLx800 microplate reader (Biotek^®^). Untreated-enzyme reactions were included in each experiment as the 100% deacetylase activity control. For each tested concentration, a corresponding blank (without HDAC6) was included to detect self-fluorescence artefacts. IC_50_ values were determined through titration with the tested compounds and data were fit using a concentration response model with GraphPad^®^ Prism 7.0.

A mass spectrometry-based assay was used to evaluate the deacetylase activity of the HDAC6-EGFP fusion proteins in lysates of HEK 293 cells. Briefly, 10.5 µM of the HDAC6 substrate BATCP (Sigma-Aldrich) was added to the diluted lysates (final HDAC6-concentrations of 50, 25, and 12.5 nM). Reactions were incubated at 37 °C for 4 h and then stopped by the addition of cold acetonitrile (2x reaction volume). The samples were kept at −80°C for 10 min and then centrifuged at 5000 x *g* for 10 min. Supernatants were analyzed on a UHPLC-ESI-MS/MS system consisting of an Acquity UPLC System (Waters, Milford, MA, USA) connected to a Quattro Micro triple quadrupole mass spectrometer equipped with an ESI source operating in positive-ion mode (Waters). Deacetylase activity was determined as previously described.^26^

### 8. ELISA-based PPI competition assay

His_6_-tagged ubiquitin was prepared at 1 µM in BupH™ Tris Buffer (Thermo Scientific, pH 7.2) and immobilized on Pierce^®^ Nickel-coated 96-well plates overnight (100 µl/well) at room temperature (RT). Plates were then washed 3x (wash buffer A: 25 mM Tris/HCl pH 7.2, 150 mM NaCl, 0.05% Tween-20). Recombinant HDAC6 WT (0.2 µM, 2% DMSO in HDAC6 assay buffer) was used as bound control (HDAC6, 100% binding to His_6_-Ub) while a pre-incubated mixture containing 0.1 µM HDAC6, 0.05-150 µM tag-free mono-Ub and 2% DMSO (HDAC6/tag-free Ub) was used as competition control. For test compounds, a pre-incubated mixture of HDAC6, 100 µM compound and 2% DMSO was used. Solutions of HDAC6 (for bound control and blank), HDAC6 with tag-free Ub (for competition control), or HDAC6 with compound (for test) were added to wells containing immobilized His_6_-Ub. Plates were then incubated for 1 h at RT and washed once (wash buffer B: 50 mM Tris/HCl pH 8.1, 137 mM NaCl, 1 mM MgCl2, 2.7 mM KCl, 0.05% Tween-20). Anti-HDAC6 rabbit antibody (1:400) was added to control and test wells, while half of the wells that contained HDAC6 alone were considered as blanks and received HDAC6 assay buffer instead, and the plate was incubated overnight at RT. Wells were washed three times (wash buffer B), followed by incubation with an anti-rabbit IgG-FITC secondary antibody (1:200) for 2 h at RT. Finally, wells were washed three times (wash buffer B) and 100 µL of HDAC6 assay buffer was added to all wells prior to fluorescence measurements on a FLx800 microplate reader (BioTek^®^, excitation at 485/20 nm and emission at 528/20 nm). PPI inhibition was calculated as HDAC6 fraction bound to His_6_-Ub, using the competition control (tag-free Ub) at 50 µM as the unbound reference. IC_50_ values were determined for concentration-response curves titrated with tag-free mono-Ub (competition control) and tested molecules. Competition data were plotted and fitted in GraphPad^®^ Prism 7.0.

## Acknowledgements

Authors are thankful to the Swiss National Science Foundation (P300P3_158507), Pierre Mercier Foundation (Switzerland), and NIH grant R01GM115997 (to REC) for financial support.

## Supplementary Information

**Fig. S1.**
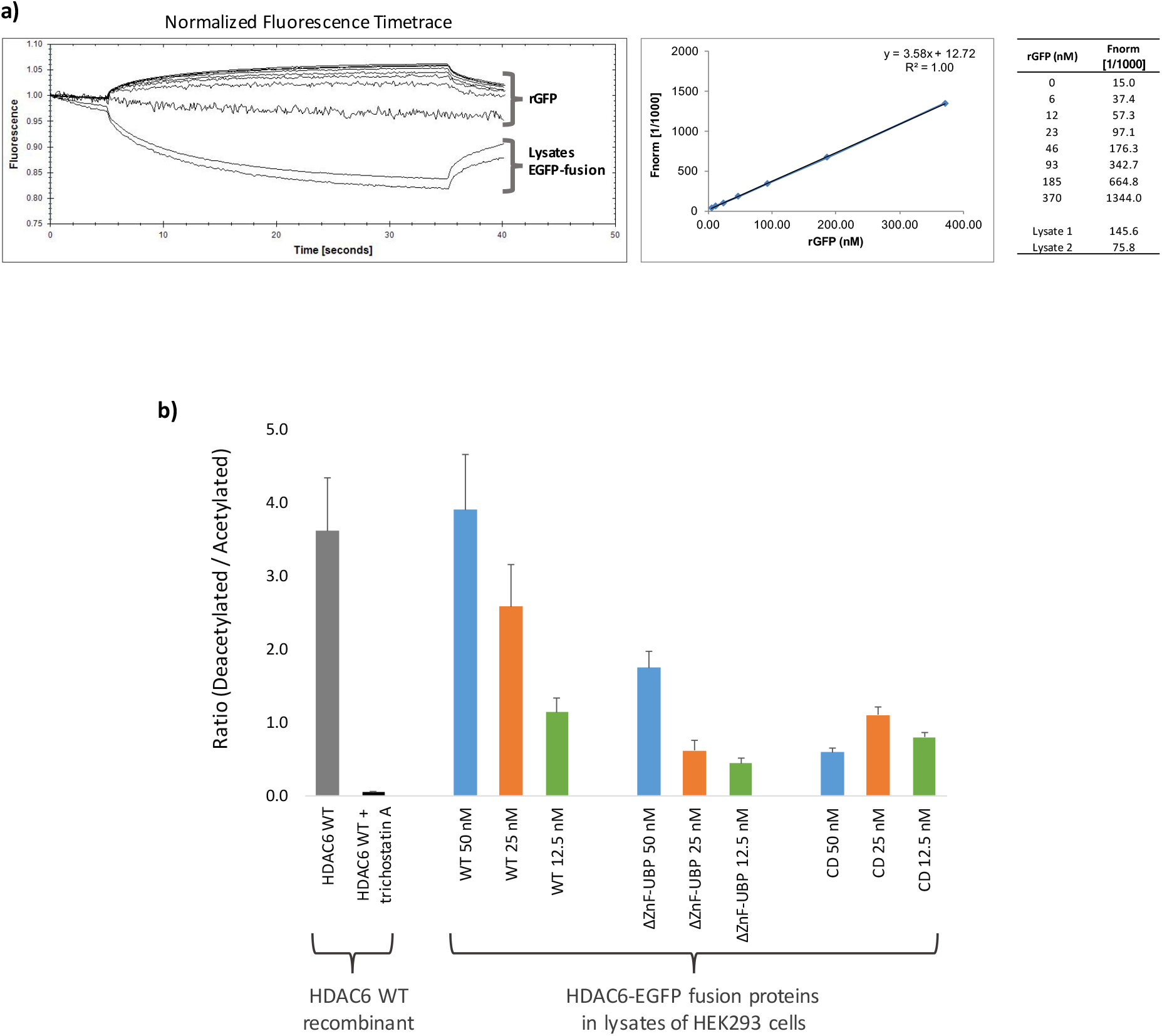
HDAC6-EGFP concentration and catalytic activity in lysates of transfected HEK 293 cells. **a)** Calibration curve for recombinant GFP (rGFP). Fluorescence of rGFP and of the EGFP-fusion proteins in the cell lysates was measured in the Monolith NT.115 Instrument (LED power 60%, excitation 460-480 nm, and emission 515-530 nm). **b)** MS-based catalytic activity of HDAC6-EGFP fusion proteins in lysates of transfected HEK 293 cells. The deacetylase activity of EGFP-fusion proteins in the lysates was measured using the HDAC6 selective substrate BATCP as previously described.^26^ Lysates were diluted to final EGFP concentrations of 50, 25, and 12.5 nM, corresponding to the concentrations of HDAC6-EGFP fusion proteins used for the MST measurements. Deacetylase activity of recombinant HDAC6 WT (50 U/reaction volume, Sigma-Aldrich) was measured as control in presence and absence of the pan-HDAC inhibitor trichostatin A (300 nM). Lysates of cells expressing HDAC6-ΔZnF-UBP showed a decrease in deacetylase activity in comparison with HDAC6 WT. On the other hand, the deacetylase activity observed for lysates of cells expressing HDAC6 CD is probably due to basal amounts of HDAC6 in HEK 293 cells enough to deacetylate the substrate, but below the detection limit by Western blot.

**Fig. S2.**
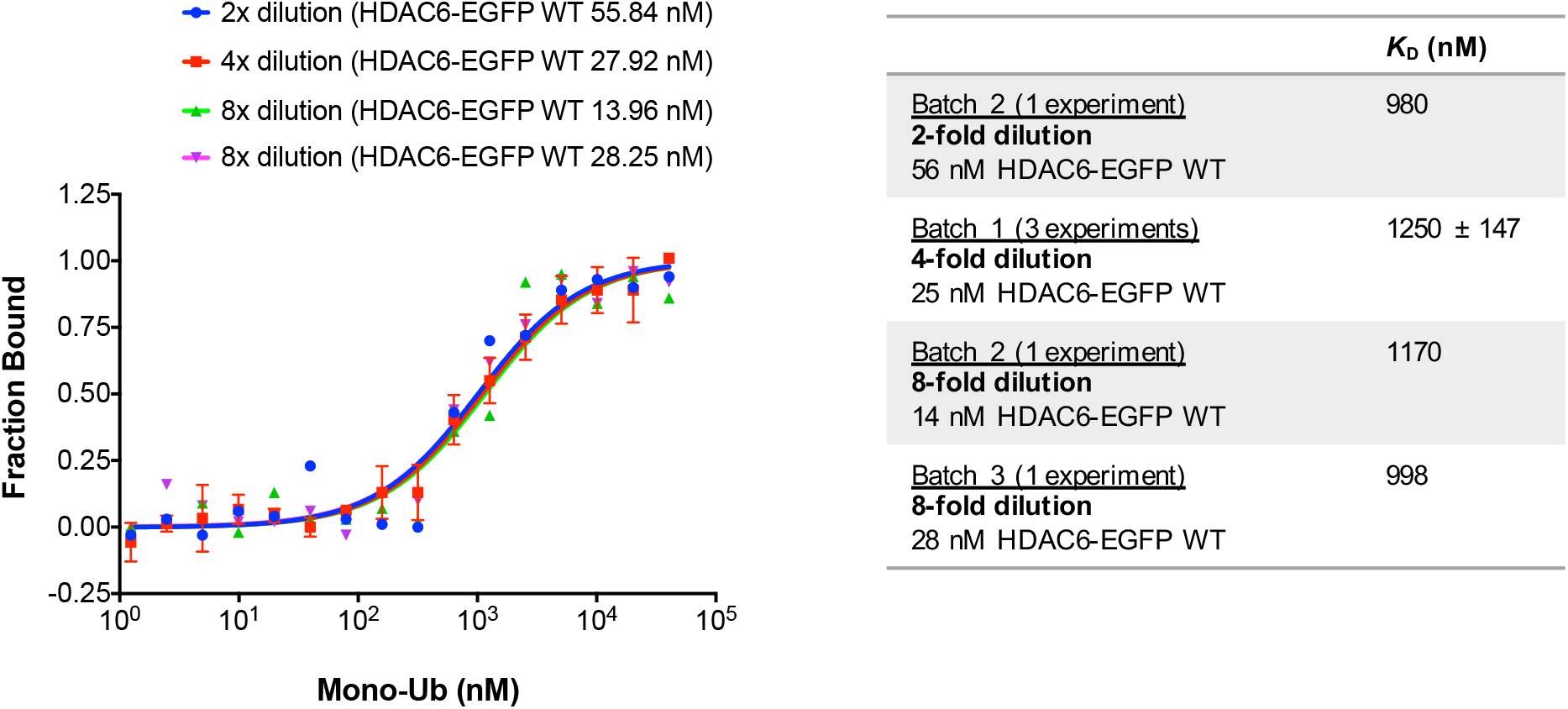
Effect of lysate dilution on MST experiments. MST experiments with different batches of lysates diluted 2-, 4- and 8-fold indicated no significant competition between recombinant mono-Ub and endogenous free Ub or other Ub-like proteins. The binding curve for batch 1 (25 nM HDAC6-EGFP WT, 4-fold dilution) is the same as presented in **Fig. 1c**.

**Fig. S3.**
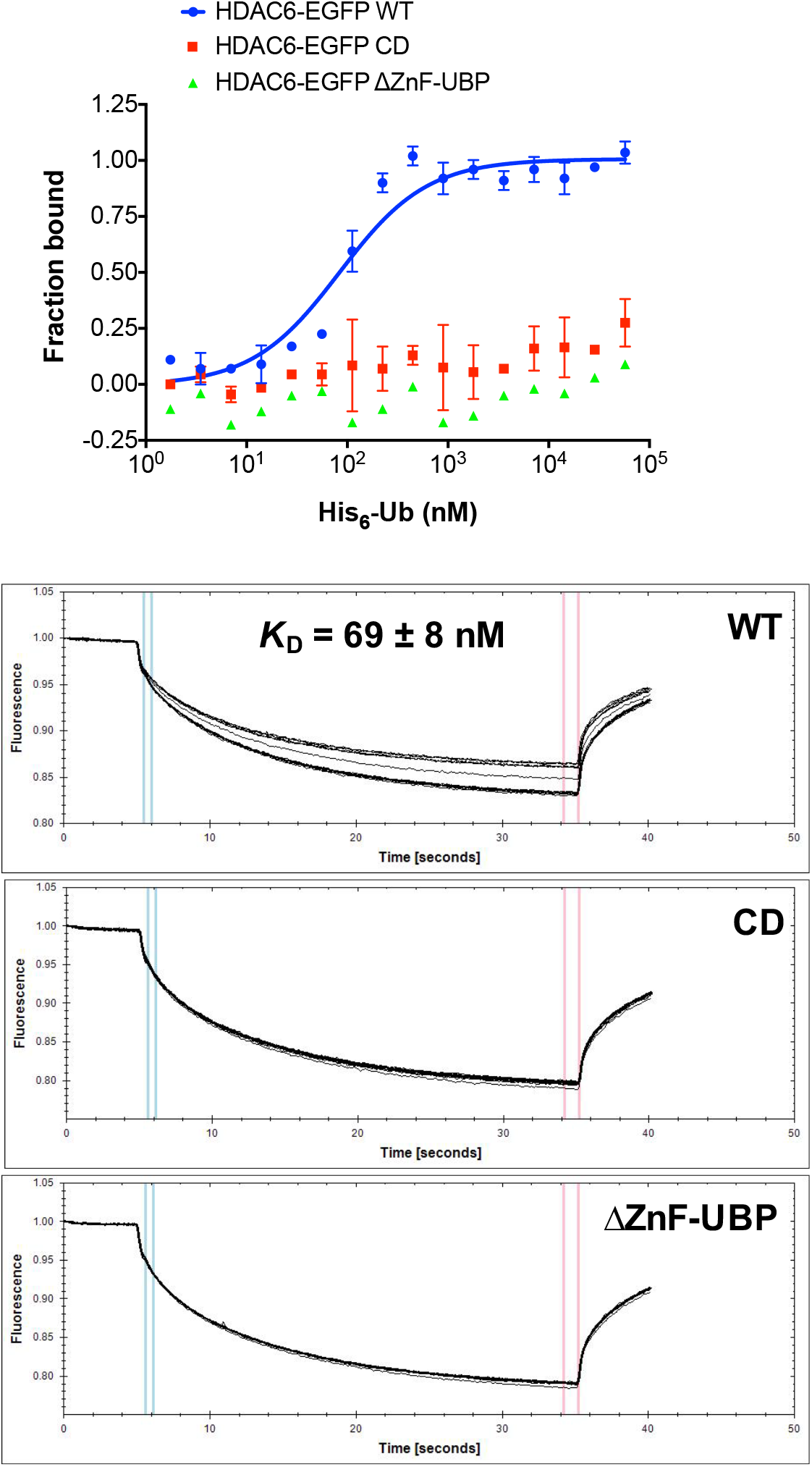
MST binding experiments with HDAC6-EGFP fusion proteins in lysates of HEK 293 cells and His_6_-ubiquitin (His_6_-Ub). Lysates of cells expressing HDAC6-EGFP WT, CD, and ΔZnF-UBP were diluted to result in EGFP final concentrations corresponding to approximately 25 nM. As observed for non-tagged mono-Ub, His_6_-Ub showed to interact with HDAC6-EGFP WT but not with HDAC6-EGFP CD or HDAC6 EGFP-ΔZnF-UBP. The *K*_D_ of 69 nM indicates that His_6_-Ub binds to HDAC6 with approximately 20-fold higher affinity than non-tagged mono-Ub. This binding affinity is close to the *K*_D_ reported for the binding between HDAC6 ZnF-UBP and His_6_-Ub (*K*_D_ = 60 nM, measured by isothermal titration calorimetry). ^27^

**Fig. S4.**
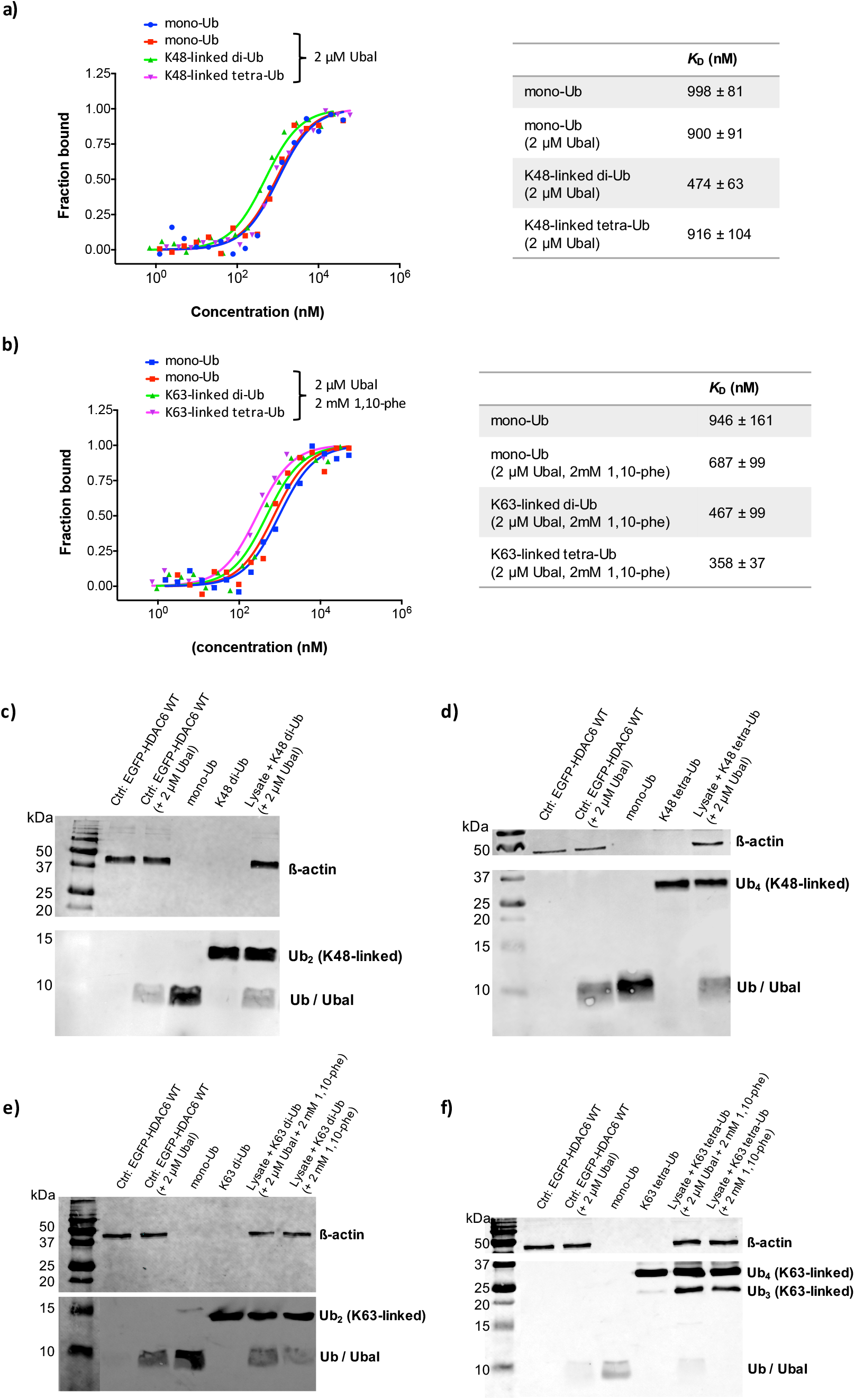
HDAC6-EGFP WT binds to mono-Ub, K48-, and K63-linked polyubiquitin chains with similar affinities. To avoid degradation by endogenous deubiquitinating enzymes (DUBs), lysates were pre-incubated with 2 µM ubiquitin aldehyde (Ubal) for experiments with K48-linked polyubiquitin chains and with 2 µM Ubal + 2 mM 1,10-phenanthroline (1,10-phe) for experiments with K63-polyubiquitin chains.^28^ **a)** MST data for mono-Ub and K48-linked polyubiquitin chains binding to HDAC6-EGFP WT. **b)** MST data for mono-Ub and K63-linked polyubiquitin chains binding to EGFP-HDAC6 WT. The observed 2-fold higher affinity measured for K48-and K63-linked di-Ub as well as the 4-fold higher affinity measured for K63-linked tetra-Ub may indicate some DUB activity in the lysates. To address this possibility, we verified the integrity of K48-and K63-linked polyubiquitin after their incubation with lysates of cells expressing the HDAC6-EGFP fusion proteins under the same conditions used in the binding assays. Lysates pre-incubated with 2 µM Ubal (15 min, 37 ºC) were further incubated with **c)** 10 µM K48-linked di-Ub (10 min, 37 °C) or **d)** 10 µM K48-linked tetra-Ub (10 min, 37 °C), and chain integrity was analyzed by Western blot. Lysates pre-incubated with 2 µM Ubal + 2 mM 1,10-phenanthroline or with 1,10-phenanthroline alone (15 min, 37 ºC) were incubated with **e)** 10 µM K63-linked di-Ub (10 min, 37 °C) or **f)** 10 µM K63-linked tetra-Ub (10 min, 37 °C) and analyzed by Western blot. Control experiments were performed with 10 µM mono-Ub or polyubiquitin chains incubated with lysis buffer. These data showed that polyubiquitin chains can be degraded by DUBs present in the lysates even after a pre-incubation step with DUB inhibitors, indicating that DUB inhibitors composition and concentration has to be carefully adjusted when the method is intended to determine affinity towards Ub-chains.

**Fig. S5.**
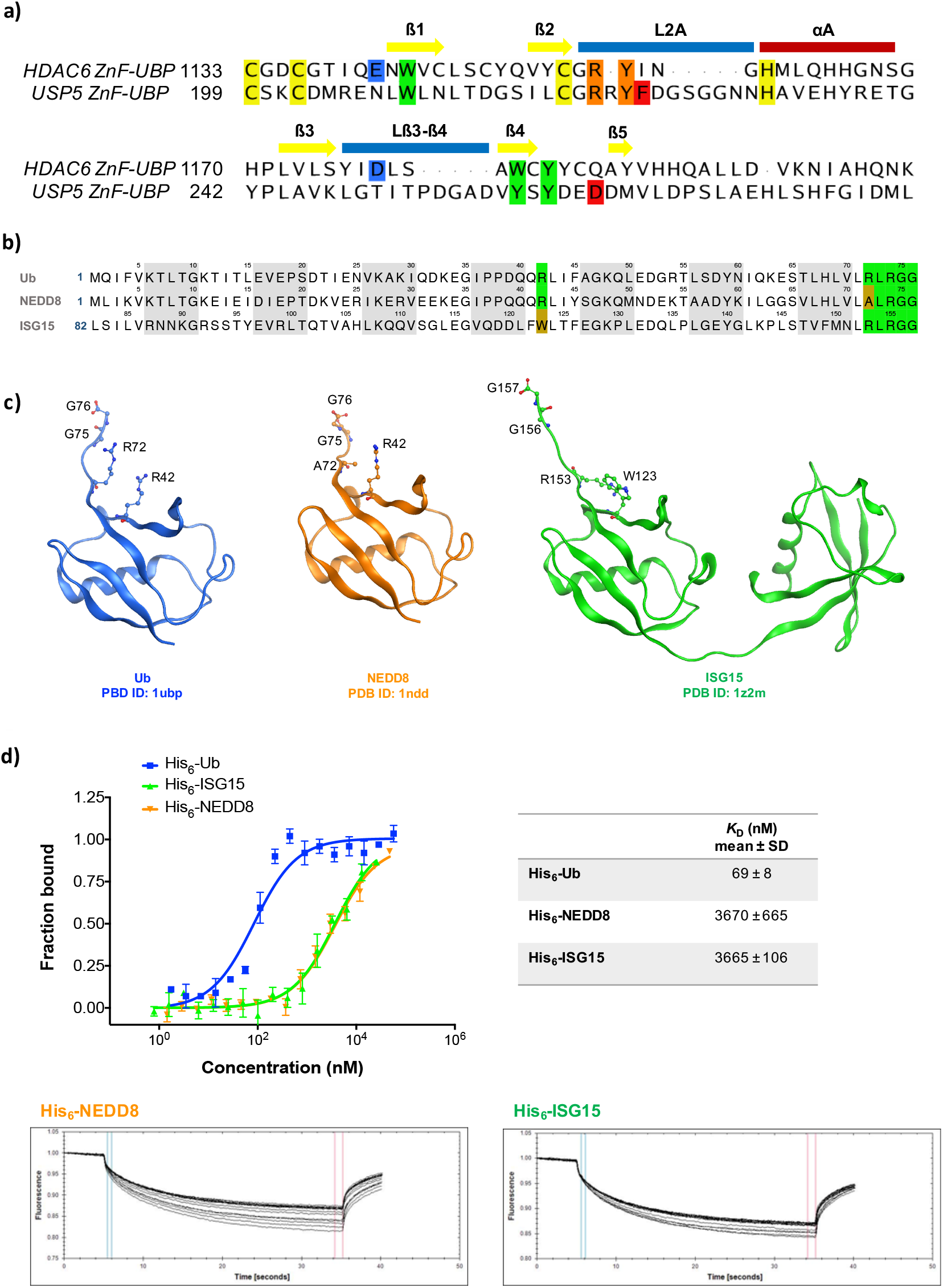
Mono-Ub binds to HDAC6-EGFP WT with 50-fold higher affinity than the Ub-like proteins NEDD8 and ISG15. **a)** Sequences of HDAC6 ZnF-UBP and USP5 ZnF-UBP. Residues coordinating with Zn^2+^ are highlighted in yellow and residues interacting with Ub are highlighted in green (aromatic pocket), orange (loop L2A), blue (HDAC6 residues non-conserved in USP5), and red (USP5 residues non-conserved in HDAC6)^4^. **b)** Sequence alignment of human Ub (Uniprot ID: P0GC48, residues 1-76) with the mature forms of human NEDD8 (Uniprot ID: Q15843, residues 1-76) and ISG15 (Uniprot ID: P05161, residues 82-157). Conserved residues contributing to HDAC6-Ub binding are highlighted in green. Non-conserved NEDD8 A72 and ISG15 W123 residues are highlighted in dark yellow. **c)** 3D structures of human Ub, NEDD8, and ISG15. **d)** MST data for His_6_-NEDD8 and His_6_-ISG15 binding to HDAC6-EGFP WT. Curve for His_6_-Ub binding to HDAC6-EGFP is the same presented in **Fig. S3**. The decreased affinity observed for NEDD8 and ISG15, when compared to mono-Ub, supports our previous molecular dynamics data suggesting that Ub R42 (replaced by a tryptophan in ISG15) and R72 (replaced by an alanine NEDD8) may contribute to HDAC6-Ub binding by making salt bridges with HDAC6 E1141 and D1178. In addition, we observed that neither mono-Ub nor the Ub-like proteins NEDD8 and ISG15 could bind HDAC6-EGFP ΔZnF-UBP (data not shown), confirming that the ZnF-UBP domain is essential for the interaction of these Ub-like proteins with HDAC6.

**Fig. S6.**
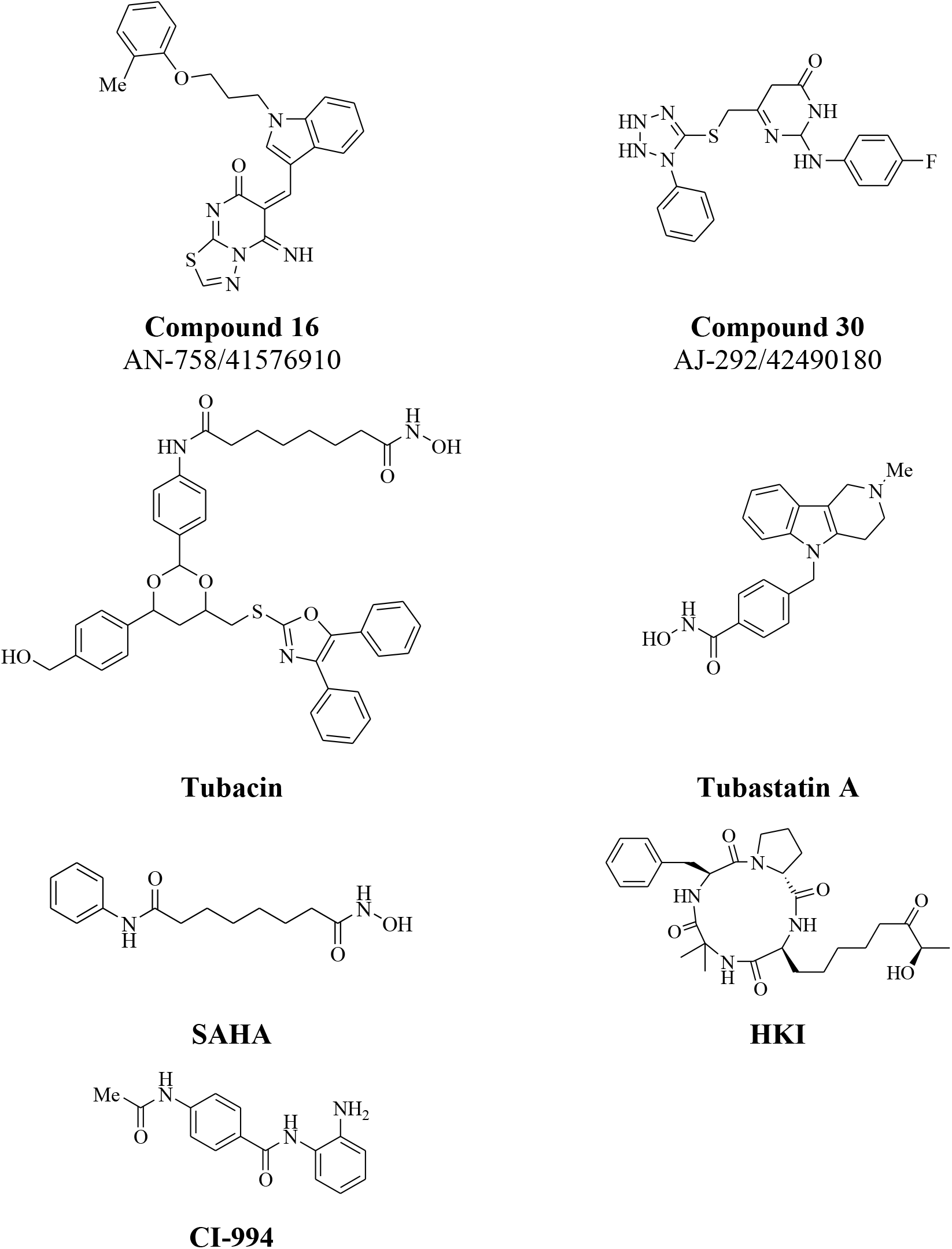
2D structures of biologically active compounds (16 and 30) selected from SBVS and positive/negative controls. Compound **16** was a hit at 100 μM in the MST screening assay for binding full-length HDAC6 and was able to inhibit HDAC6-Ub PPI in the ELISA-based assay (IC_50_=78 μM). Compound **30** was shown to inhibit the HDAC6-Ub PPI in the ELISA-based assay (IC_50_=95 μM). Tubacin and tubastatin A are known catalytic inhibitors selective to HDAC6. HKI and CI-994 are selective inhibitors of class I (nuclear) HDACs.

**Fig. S7.**
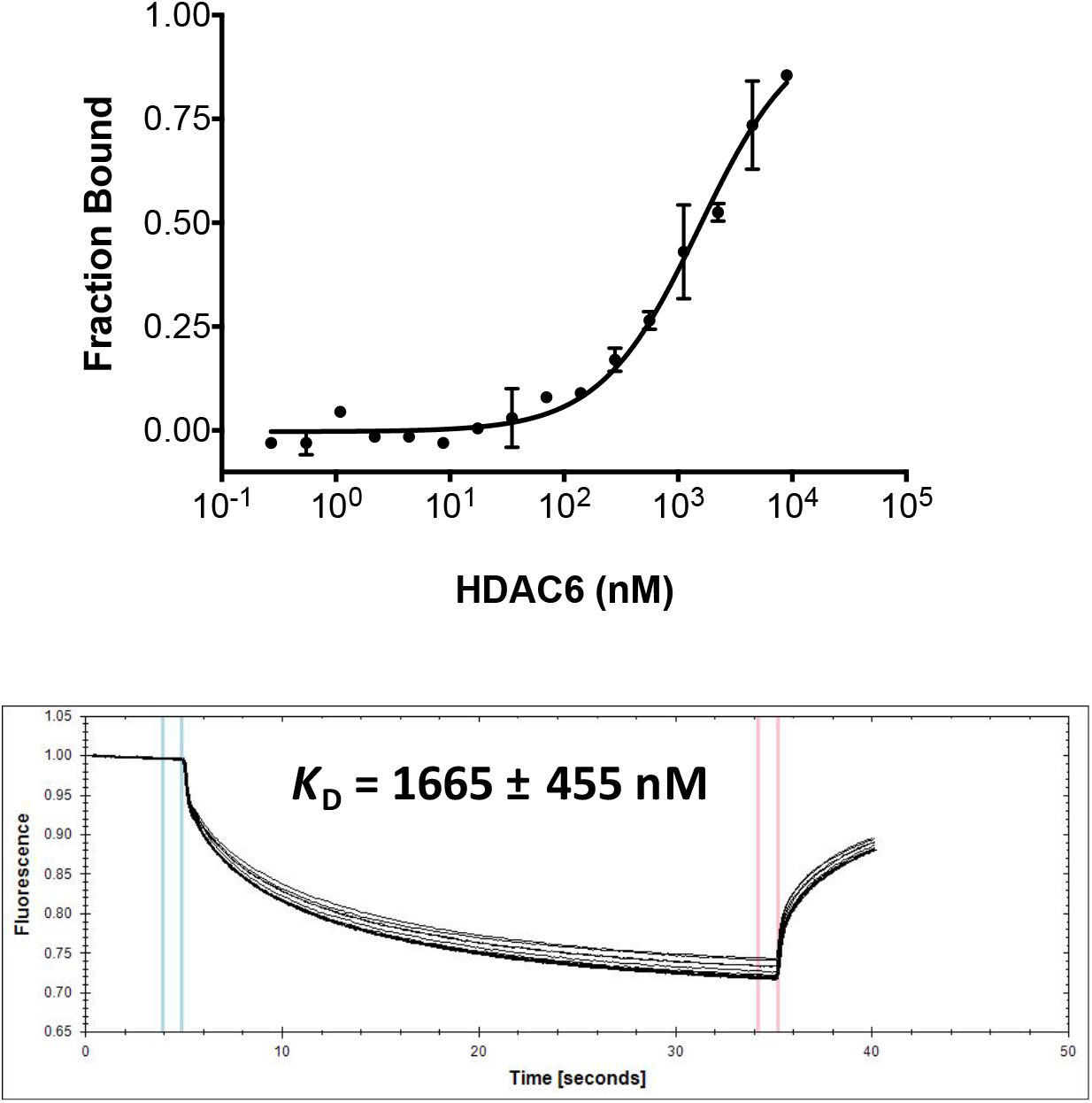
Binding of recombinant full-length HDAC6 WT to mono-Ub labeled with NT-647 using NHS coupling chemistry. Mono-Ub labeled with the fluorescent dye NT-647 (66 nM final concentrations) was titrated with recombinant HDAC6 WT (0.3 – 9000 nM).

**Table S1.**
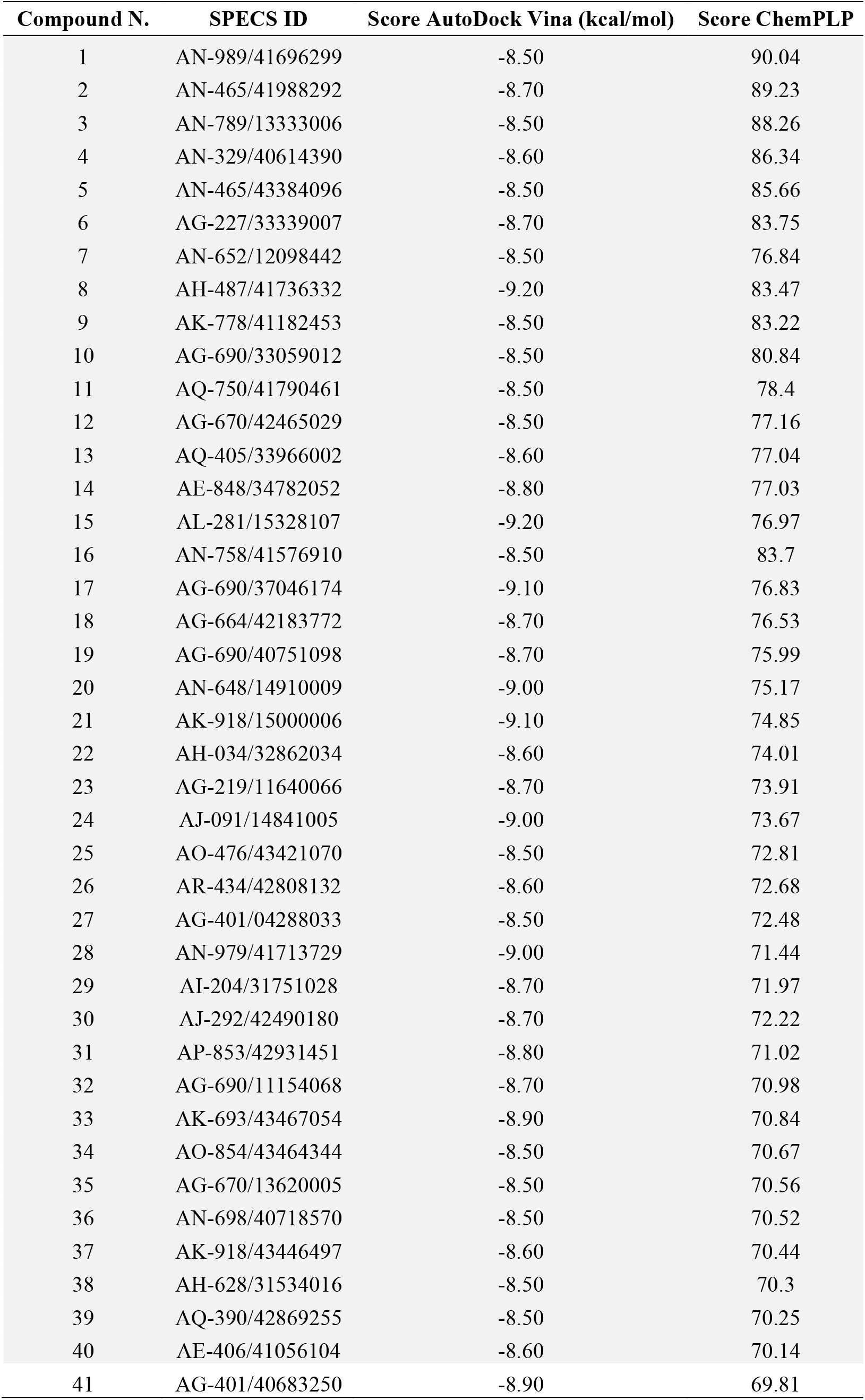
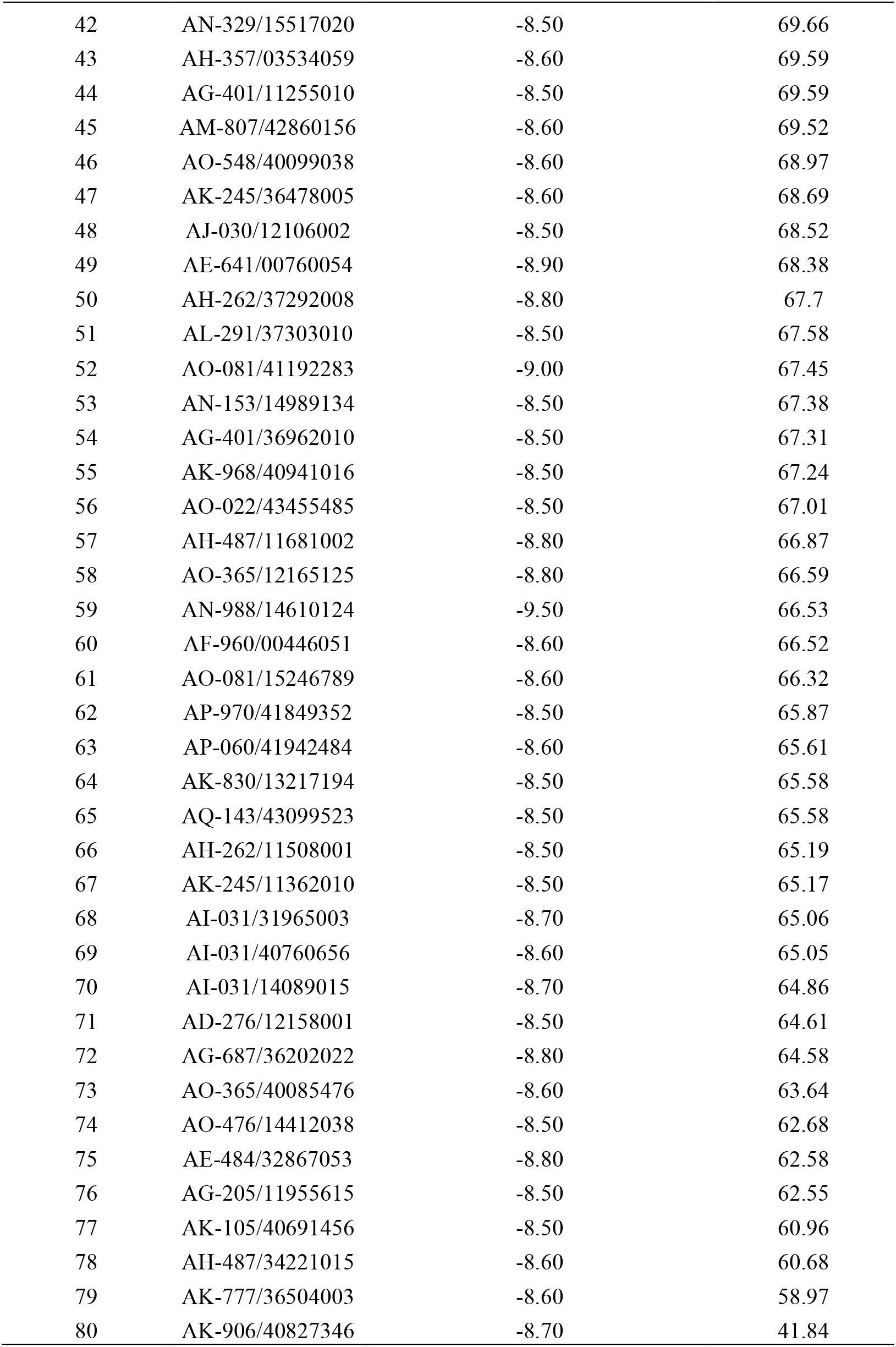
Summary of the SBVS results obtained by using Autodock Vina and GOLD molecular docking approaches to screen the Specs database against HDAC6 ZnF-UBP. 80 compounds with a Vina score lower than −8.5 kcal/mol were selected by visual inspection. Redocking experiments with GOLD allowed the selection of 40 compounds for *in vitro* studies, possessing ChemPLP scores higher than 70, and here shaded in gray.

| Compound N. | SPECS ID | Score AutoDock Vina (kcal/mol) | Score ChemPLP |
| --- | --- | --- | --- |
| 1 | AN-989/41696299 | -8.50 | 90.04 |
| 2 | AN-465/41988292 | -8.70 | 89.23 |
| 3 | AN-789/13333006 | -8.50 | 88.26 |
| 4 | AN-329/40614390 | -8.60 | 86.34 |
| 5 | AN-465/43384096 | -8.50 | 85.66 |
| 6 | AG-227/33339007 | -8.70 | 83.75 |
| 7 | AN-652/12098442 | -8.50 | 76.84 |
| 8 | AH-487/41736332 | -9.20 | 83.47 |
| 9 | AK-778/41182453 | -8.50 | 83.22 |
| 10 | AG-690/33059012 | -8.50 | 80.84 |
| 11 | AQ-750/41790461 | -8.50 | 78.4 |
| 12 | AG-670/42465029 | -8.50 | 77.16 |
| 13 | AQ-405/33966002 | -8.60 | 77.04 |
| 14 | AE-848/34782052 | -8.80 | 77.03 |
| 15 | AL-281/15328107 | -9.20 | 76.97 |
| 16 | AN-758/41576910 | -8.50 | 83.7 |
| 17 | AG-690/37046174 | -9.10 | 76.83 |
| 18 | AG-664/42183772 | -8.70 | 76.53 |
| 19 | AG-690/40751098 | -8.70 | 75.99 |
| 20 | AN-648/14910009 | -9.00 | 75.17 |
| 21 | AK-918/15000006 | -9.10 | 74.85 |
| 22 | AH-034/32862034 | -8.60 | 74.01 |
| 23 | AG-219/11640066 | -8.70 | 73.91 |
| 24 | AJ-091/14841005 | -9.00 | 73.67 |
| 25 | AO-476/43421070 | -8.50 | 72.81 |
| 26 | AR-434/42808132 | -8.60 | 72.68 |
| 27 | AG-401/04288033 | -8.50 | 72.48 |
| 28 | AN-979/41713729 | -9.00 | 71.44 |
| 29 | AI-204/31751028 | -8.70 | 71.97 |
| 30 | AJ-292/42490180 | -8.70 | 72.22 |
| 31 | AP-853/42931451 | -8.80 | 71.02 |
| 32 | AG-690/11154068 | -8.70 | 70.98 |
| 33 | AK-693/43467054 | -8.90 | 70.84 |
| 34 | AO-854/43464344 | -8.50 | 70.67 |
| 35 | AG-670/13620005 | -8.50 | 70.56 |
| 36 | AN-698/40718570 | -8.50 | 70.52 |
| 37 | AK-918/43446497 | -8.60 | 70.44 |
| 38 | AH-628/31534016 | -8.50 | 70.3 |
| 39 | AQ-390/42869255 | -8.50 | 70.25 |
| 40 | AE-406/41056104 | -8.60 | 70.14 |
| 41 | AG-401/40683250 | -8.90 | 69.81 |
| 42 | AN-329/15517020 | -8.50 | 69.66 |
| 43 | AH-357/03534059 | -8.60 | 69.59 |
| 44 | AG-401/11255010 | -8.50 | 69.59 |
| 45 | AM-807/42860156 | -8.60 | 69.52 |
| 46 | AO-548/40099038 | -8.60 | 68.97 |
| 47 | AK-245/36478005 | -8.60 | 68.69 |
| 48 | AJ-030/12106002 | -8.50 | 68.52 |
| 49 | AE-641/00760054 | -8.90 | 68.38 |
| 50 | AH-262/37292008 | -8.80 | 67.7 |
| 51 | AL-291/37303010 | -8.50 | 67.58 |
| 52 | AO-081/41192283 | -9.00 | 67.45 |
| 53 | AN-153/14989134 | -8.50 | 67.38 |
| 54 | AG-401/36962010 | -8.50 | 67.31 |
| 55 | AK-968/40941016 | -8.50 | 67.24 |
| 56 | AO-022/43455485 | -8.50 | 67.01 |
| 57 | AH-487/11681002 | -8.80 | 66.87 |
| 58 | AO-365/12165125 | -8.80 | 66.59 |
| 59 | AN-988/14610124 | -9.50 | 66.53 |
| 60 | AF-960/00446051 | -8.60 | 66.52 |
| 61 | AO-081/15246789 | -8.60 | 66.32 |
| 62 | AP-970/41849352 | -8.50 | 65.87 |
| 63 | AP-060/41942484 | -8.60 | 65.61 |
| 64 | AK-830/13217194 | -8.50 | 65.58 |
| 65 | AQ-143/43099523 | -8.50 | 65.58 |
| 66 | AH-262/11508001 | -8.50 | 65.19 |
| 67 | AK-245/11362010 | -8.50 | 65.17 |
| 68 | AI-031/31965003 | -8.70 | 65.06 |
| 69 | AI-031/40760656 | -8.60 | 65.05 |
| 70 | AI-031/14089015 | -8.70 | 64.86 |
| 71 | AD-276/12158001 | -8.50 | 64.61 |
| 72 | AG-687/36202022 | -8.80 | 64.58 |
| 73 | AO-365/40085476 | -8.60 | 63.64 |
| 74 | AO-476/14412038 | -8.50 | 62.68 |
| 75 | AE-484/32867053 | -8.80 | 62.58 |
| 76 | AG-205/11955615 | -8.50 | 62.55 |
| 77 | AK-105/40691456 | -8.50 | 60.96 |
| 78 | AH-487/34221015 | -8.60 | 60.68 |
| 79 | AK-777/36504003 | -8.60 | 58.97 |
| 80 | AK-906/40827346 | -8.70 | 41.84 |

**Table S2.**
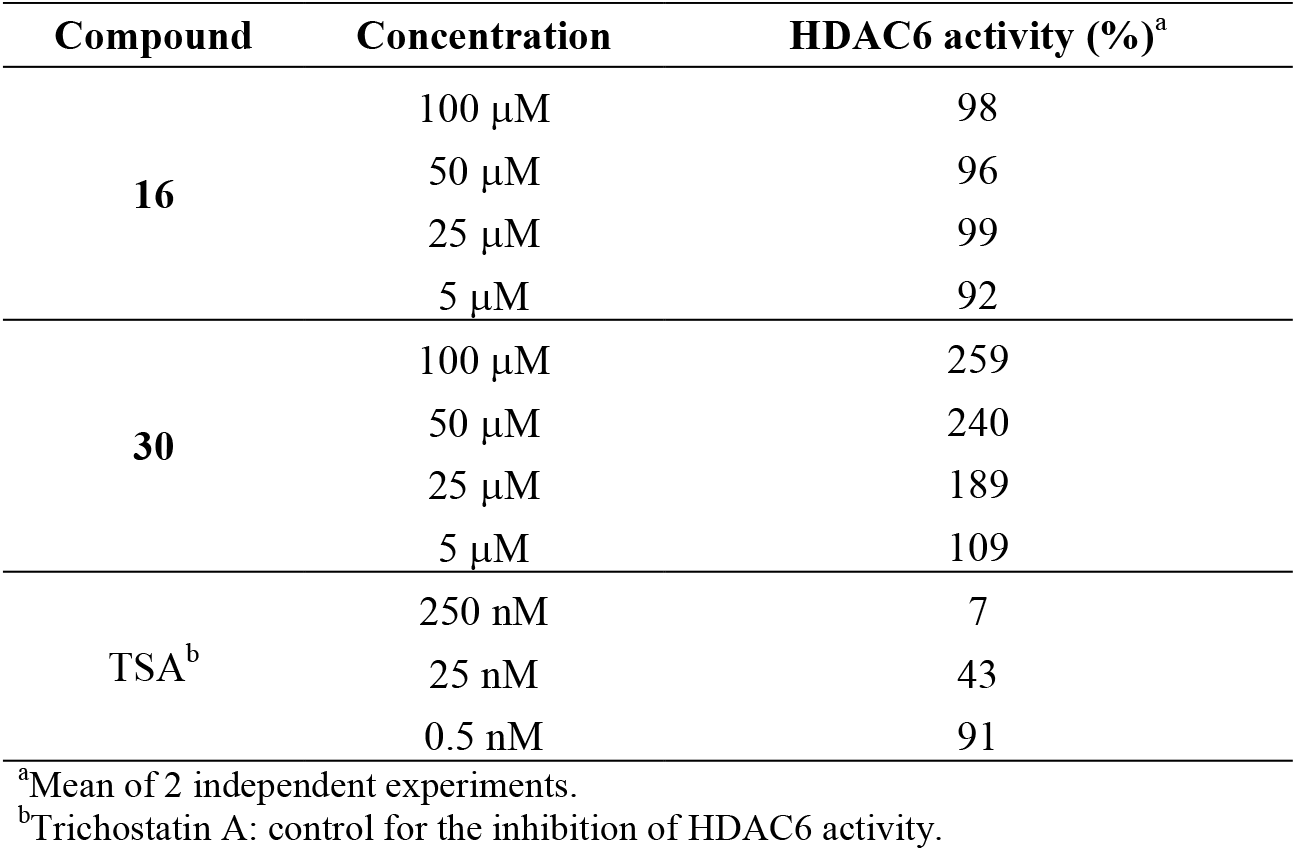
Effect of selected hits on HDAC6 catalytic activity.

| Compound | Concentration | HDAC6 activity (%)<sup>a</sup> |
| --- | --- | --- |
| **16** | 100 μM | 98 |
| | 50 μM | 96 |
| | 25 μM | 99 |
| | 5 μM | 92 |
| **30** | 100 μM | 259 |
| | 50 μM | 240 |
| | 25 μM | 189 |
| | 5 μM | 109 |
| TSA<sup>b</sup> | 250 nM | 7 |
|  | 25 nM | 43 |
|  | 0.5 nM | 91 |

<sup>a</sup>Mean of 2 independent experiments.<sup>b</sup>Trichostatin A: control for the inhibition of HDAC6 activity.

